# Zinc protection of fertilized eggs is an ancient feature of sexual reproduction in animals

**DOI:** 10.1101/2020.06.08.140541

**Authors:** Katherine L. Wozniak, Rachel E. Bainbridge, Dominique W. Summerville, Maiwase Tembo, Wesley A. Phelps, Monica L. Sauer, Bennett W. Wisner, Madelyn E. Czekalski, Srikavya Pasumarthy, Meghan L. Hanson, Melania B. Linderman, Catherine H. Luu, Madison E. Boehm, Steven M. Sanders, Katherine M. Buckley, Daniel J. Bain, Matthew L. Nicotra, Miler T. Lee, Anne E. Carlson

## Abstract

One of the earliest and most prevalent barriers to successful reproduction is polyspermy, or fertilization of an egg by multiple sperm. To prevent these supernumerary fertilizations, eggs have evolved multiple mechanisms. It has recently been proposed that zinc released by mammalian eggs at fertilization may block additional sperm from entering. Here, we demonstrate that eggs from amphibia and teleost fish also release zinc. Using *Xenopus laevis* as a model, we document that zinc reversibly blocks fertilization. Finally, we demonstrate that extracellular zinc similarly disrupts early embryonic development in eggs from diverse phyla, including: Cnidaria, Echinodermata, and Chordata. Our study reveals that a fundamental strategy protecting human eggs from fertilization by multiple sperm may have evolved more than 650 million years ago.

## Introduction

Fertilization of an egg by more than one sperm, a condition known as polyspermy, is lethal for most animals. Consequently, eggs use several strategies to shield nascent zygotes from penetration by additional sperm [1]; the two most common are referred to as the *fast* and *slow blocks to polyspermy*. In the fast block, fertilization immediately changes the electrical charge of the egg’s membrane [2–4]. This shift to a positive potential is sufficient to prevent sperm entry [5], but is only used by externally fertilized eggs. By contrast, eggs from most sexual reproducers, including mammals, use the slow block [1, 6]. During the slow block to polyspermy, an increase of intracellular calcium causes eggs to release materials that form an extracellular barrier impenetrable to sperm [7, 8]. The slow block initiates at least three processes that prevent future fertilizations: a membrane-modification that stops sperm binding [7, 9], and two changes to the extracellular matrix (called the zona pellucida in mammals) that block sperm binding and penetration [8, 10]. We are just beginning to understand how the released materials cause these transformations to protect the nascent zygote.

Recently, extracellular zinc has been proposed to keep fertilized eggs from polyspermy as part of the slow block [11, 12]. Zinc release during the slow block has been documented in mammalian eggs from humans [13], mice [14], and cows [15]. Zinc release has also been linked to gamete maturation, cell cycle resumption, and initiation of embryonic development [14, 16–18]. We sought to determine the conservation of this zinc release in diverse species with varied fertilization strategies, and more directly probe whether extracellular zinc protects eggs from additional fertilizations. We assayed for a role of zinc release from eggs of the African clawed frog *Xenopus laevis*, which are external fertilizers; both the fast and slow polyspermy blocks have been extensively characterized in this model organism [1]. We also used a salamander, the axolotl (*Ambystoma mexicanum*). Salamanders and frogs are both amphibians with eggs that employ the slow polyspermy block, yet salamanders are internal fertilizers whose eggs do not depolarize with sperm entry [19]. Zinc release was also observed from zebrafish (*Danio rerio*) eggs; unlike mammalian and amphibian eggs, zebrafish eggs activate upon hydration in a fertilization-independent process [20]. Finally, we used two invertebrate species with highly tractable fertilization from distantly related phyla: the purple sea urchin *Strongylocentrotus purpuratus* (Echinodermata), and the hydroid *Hydractinia symbiolongicarpus* (Cnideria). Probing fertilization and activation in eggs from these five species, we now demonstrate that the zinc release from activated eggs is shared by distantly related animals and that extracellular zinc-protection of egg from supernumerary fertilizations is an ancient phenomenon.

## Results and discussion

To determine whether *X. laevis* eggs release zinc upon fertilization, we used confocal microscopy with the cell-impermeant, fluorescent zinc indicator FluoZin-3, before and after insemination. We observed that *X. laevis* eggs were enveloped by a singular wave of extracellular zinc within minutes of sperm addition (Fig 1A-B, S1A Fig, Movie 1). This zinc spark was observed in 16 of 20 imaged eggs. In mammalian eggs, zinc is exocytosed from cortical granules during the slow block to polyspermy [21, 22]. If zinc release also occurred during the slow block in *X. laevis* eggs, we predicted that the appearance of extracellular zinc would share characteristics, and coincide, with markers of the slow block. The observed appearance of a single zinc wave was expected because *X. laevis* eggs have one intracellular calcium transient responsible for a singular cortical granule exocytosis [23]. Following sperm entry, the envelope separates from the egg when proteases from the cortical granules cleave contacts between these two structures [1, 24]. Appearance and localization of extracellular zinc was in sync with the lifting of the envelope (Fig 1A *red arrows*). Cortical contraction was observed in all imaged eggs (Fig 1C), indicating successful fertilization and activation [25]. Zinc release was no synchronous between proximal eggs; when 2 eggs were simultaneously imaged (Movie 1), the second egg began zinc release 105 seconds after the first. We expect this is due to a difference in fertilization timing.

**Figure 1.**
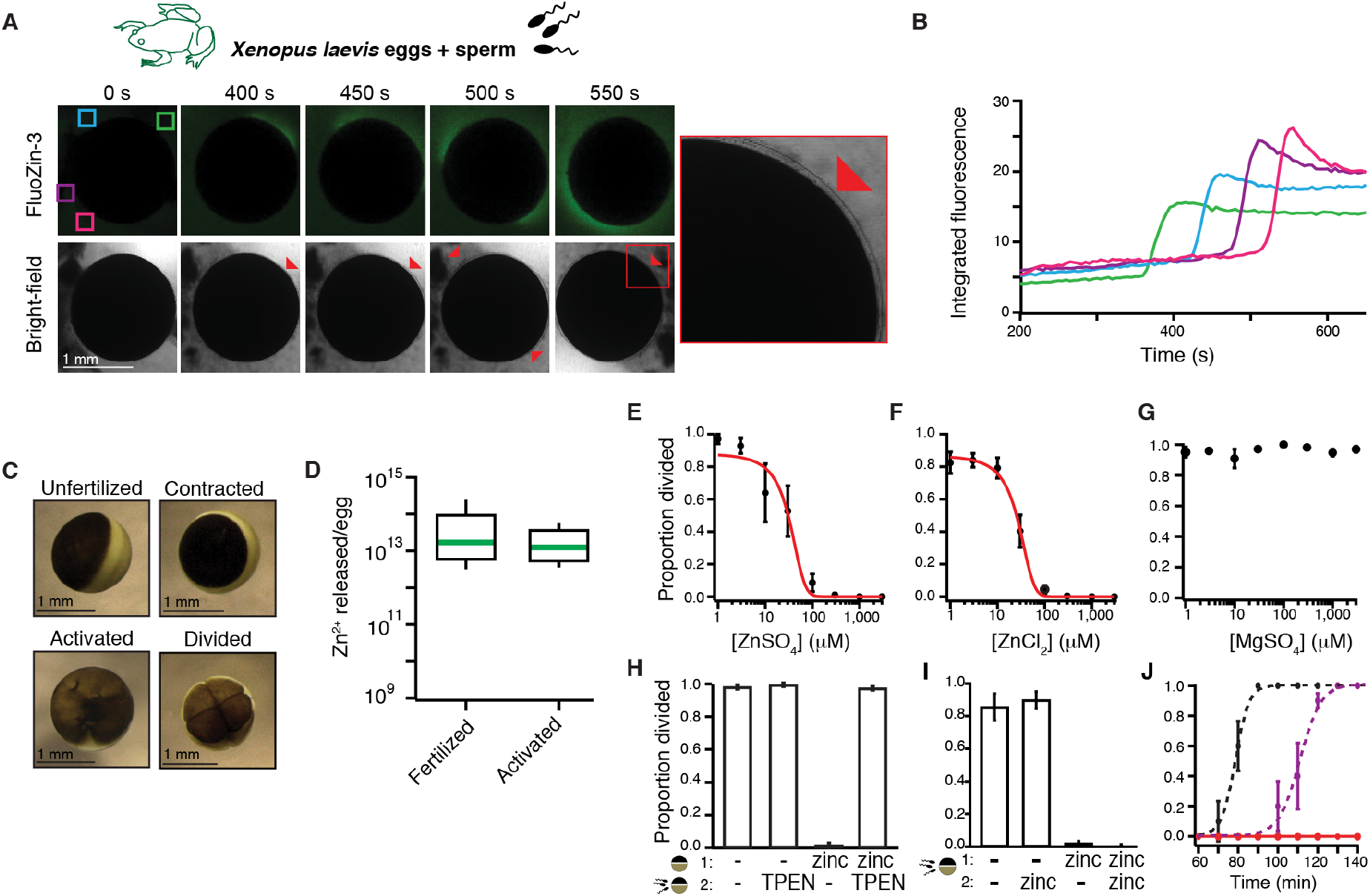
*X. laevis* eggs release zinc at fertilization, which protects eggs from additional fertilizations. (A) Representative fluorescence and bright-field images of *X. laevis* eggs in FluoZin-3, before and after sperm application. Time is relative to sperm addition. Zinc release coincided with lifting of the envelope (red arrowheads and insert) (N=16 eggs, 4 trials). (B) Integrated fluorescence, relative to time of sperm addition, detected by region of interest analysis, indicated by colored boxes in panel A. (C) Representative images of *X. laevis* eggs and embryos. (D) Box plot distribution of zinc ions released per *X. laevis* egg upon fertilization or activation with 10 μM ionomycin as detected by FluoZin-3 fluorometry. (E-G) Proportion of inseminated eggs that developed cleavage furrows in indicated concentrations of ZnSO_4_, ZnCl_2_, or MgSO_4_. Plots in A and B were fit with sigmoidal functions. (H) Proportion of development of eggs subjected to a 15 min pre-treatment with 0 or 300 μM ZnSO_4_ (solution *1*) then washed and moved to solution (*2*) with 0 or 300 μM TPEN for sperm addition (N=68-259 eggs, 5 trials). (I) Incidence of cleavage furrow development from eggs inseminated in 0 or 1 mM ZnSO_4_ then transferred to a new solution with 0 or 1 mM ZnSO_4_, 30 min after sperm addition (N=52-258 eggs in 6 trials). (J) Proportion of cleavage furrow development from eggs inseminated in and transferred to control conditions (black), inseminated in 300 μM ZnSO_4_ and transferred to 600 μM TPEN 30 min following sperm addition (purple), or eggs inseminated in and transferred to 1 mM ZnSO_4_ (red) (N=26-58 eggs in 7 trials). (E-J) Errors are s.e.m.

We explored whether signaling the slow block without fertilization would initiate zinc release from *X. laevis* eggs. Extracellular zinc was imaged before and after application of the calcium ionophore ionomycin [26], which increases cytosolic calcium to evoke cortical granule release. In both eggs and *in vitro* matured oocytes, a zinc release appeared with ionomycin application (S1A-E Fig) and also coincided with lifting of the envelope (Movie 2). Envelope lifting and cortical contraction verified successful activation (Fig 1C).

To confirm that the change in FluoZin-3 fluorescence was due to increased extracellular zinc, we fertilized *X. laevis* eggs in the zinc chelator N,N,N’,N’-tetrakis(2-pyridinylmethyl)-1,2-ethanediamine (TPEN) [27]. Under these conditions, fertilization did not evoke increased FluoZin-3 fluorescence (S1F Fig, Movie 3). Yet envelope lifting was observed in TPEN, as the chelation of zinc did not affect the cortical granule exocytosis that leads to envelope lifting. Finally, we queried proteomic [28] and RNA-sequencing [29] datasets to verify that *X. laevis* eggs express zinc transporters needed to acquire and regulate the metal. Indeed, both ZnTs and ZIPs are present in *X. laevis* eggs (S2 Fig).

Zinc released into the surrounding solution was quantified with FluoZin-3 fluorometry. Fertilization and activation with ionomycin induced an average release of 5.5 ± 2.7 x 10^13^ and 1.9 ± 1.0 x 10^13^ zinc ions/egg, respectively (Fig 1D, Table 1). Inductively coupled plasma mass spectrometry (ICP-MS) substantiated an average release of 5.8 ± 7.0 x 10^12^ zinc ions per ionomycin-treated egg (Table 1). Based on atomic absorption spectroscopy, *X. laevis* eggs have an average of 6 x 10^14^ zinc ions [30], thereby revealing that each egg loses less than 10% of its total zinc during the slow block.

**Table 1.**
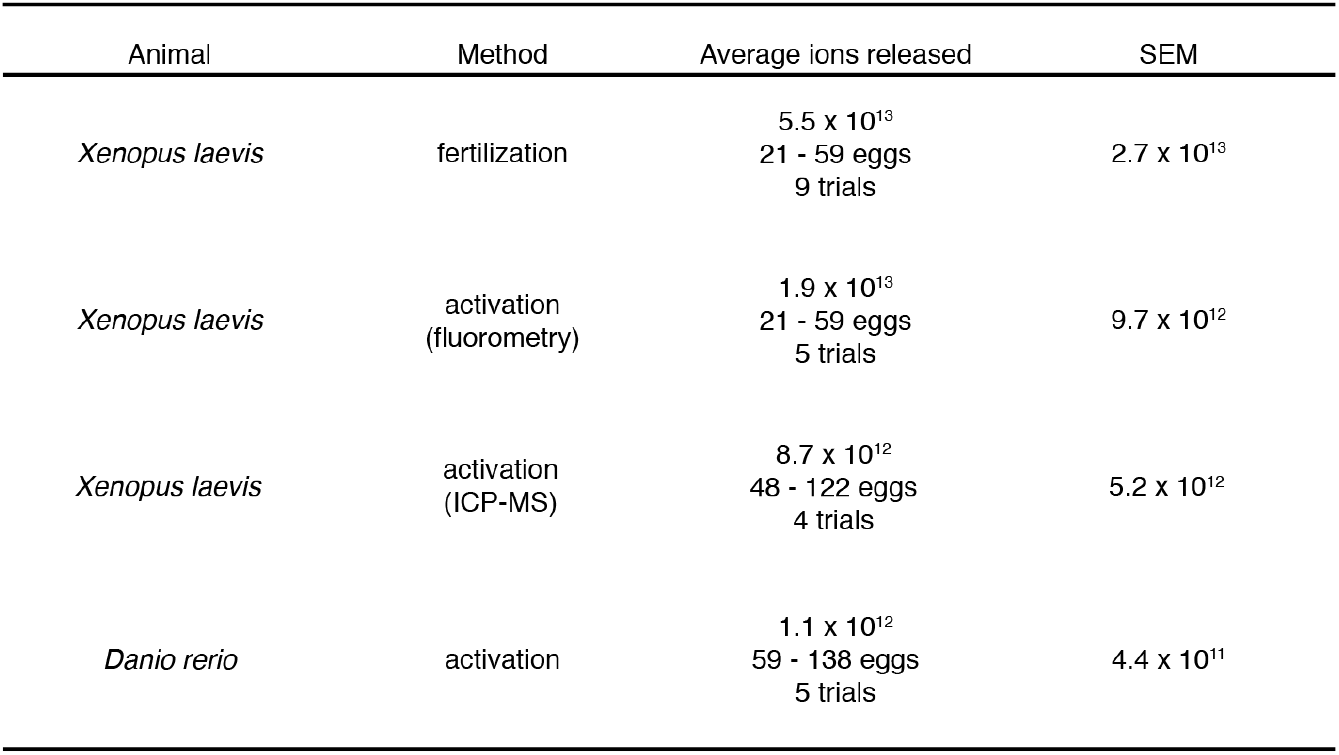
Average zinc ion release following fertilization or activation of eggs.

| Animal | Method | Average ions released | SEM |
| --- | --- | --- | --- |
| *Xenopus laevis* | fertilization | $5.5 \times 10^{13}$<br>21 - 59 eggs<br>9 trials | $2.7 \times 10^{13}$ |
| *Xenopus laevis* | activation<br>(fluorometry) | $1.9 \times 10^{13}$<br>21 - 59 eggs<br>5 trials | $9.7 \times 10^{12}$ |
| *Xenopus laevis* | activation<br>(ICP-MS) | $8.7 \times 10^{12}$<br>48 - 122 eggs<br>4 trials | $5.2 \times 10^{12}$ |
| *Danio rerio* | activation | $1.1 \times 10^{12}$<br>59 - 138 eggs<br>5 trials | $4.4 \times 10^{11}$ |

We next explored a possible role for zinc in the slow polyspermy block by inseminating *X. laevis* eggs in varying concentrations of extracellular zinc and monitoring for an indicator of early embryonic development, the appearance of cleavage furrows. If extracellular zinc protects eggs from sperm entry, we predicted that embryonic development would only occur in eggs inseminated in minimal zinc solution. We found that ZnSO_4_ inhibited the appearance of cleavage furrows in a concentration-dependent manner (Fig 1E, S1 Table); no development was observed in eggs inseminated in ≥300 μM ZnSO_4_. A sigmoidal fit of the incidence of development versus ZnSO_4_ concentration showed that half the eggs developed cleavage furrows at a zinc concentration (half-maximal inhibitory concentration, IC_50_) of 31 ± 10 μM (Fig 1E). To confirm that zinc (Zn^2+^), and not sulfate (SO_4_^2-^), was responsible for disrupting fertilization and embryonic development, *X. laevis* eggs were fertilized in varying concentrations of ZnCl_2_ or MgSO_4_ (Fig 1F-G). Whereas ZnCl_2_ inhibited development with a nearly identical concentration response to ZnSO_4_ (30 ± 8 μM), MgSO_4_ had no effect (S1 Table). To test if the effect of zinc was reversible, eggs were incubated in zinc, then transferred to and inseminated in a solution with TPEN. Nearly all eggs pretreated with zinc and subsequently inseminated in TPEN underwent cleavage (Fig 1H), thereby revealing that extracellular zinc effects on *X. laevis* eggs are reversible. In control experiments, we assayed for egg rolling, cortical contraction, or appearance of cleavage furrows to determine whether TPEN treatment without sperm was sufficient to activate *X. laevis* eggs. No signs of TPEN-induced activation were observed in any eggs (N=82 eggs in 6 independent trials).

Several critical events occur during the 90 minutes between fertilization and the appearance of cleavage furrows. If extracellular zinc prevented fertilization to thereby inhibit the appearance of cleavage furrows, we reasoned that transferring zinc-inseminated eggs to a non-zinc solution should not rescue development. Indeed, we found that *X. laevis* eggs inseminated in zinc failed to develop cleavage furrows regardless of their treatment 30 minutes after sperm addition (Fig 1I). By contrast, eggs inseminated without added zinc developed normally, even when treated with zinc 30 minutes after sperm addition (Fig 1I). This latter finding suggests that after fertilization, zinc treatment is not sufficient to halt embryonic development. Together these results reveal extracellular zinc interferes with development within 30 minutes of insemination.

To further ensure that extracellular zinc blocked fertilization, we examined how rapidly TPEN application recovered cleavage furrow appearance from the zinc-induced block of embryonic development. Eggs were fertilized with or without zinc, then 30 minutes following sperm addition, transferred to different solutions with or without TPEN. We predicted that if zinc blocked fertilization, the appearance of cleavage furrows in eggs inseminated in zinc would be shifted by the time of TPEN addition (30 minutes for these experiments). If zinc blocked another event in early embryonic development, the appearance of cleavage furrows would appear by an intermediate time. We found that under control conditions, approximately half of the eggs developed cleavage furrows 79 minutes after sperm addition (± 0.3 min, Fig 1J). By contrast, half of the eggs inseminated in zinc, then transferred to TPEN developed cleavage furrows 111 min after sperm addition (± 0.9 min), or 81 minutes following transfer to TPEN. In the presence of zinc, sperm are evidently unable to penetrate the egg. We hypothesize that sperm embedded in the jelly coat of zinc treated eggs likely fertilized upon transfer to the TPEN solution. These results indicate that zinc released from *X. laevis* eggs following fertilization can protect the zygote from fertilization by additional sperm.

We next tested for zinc release from gametes of another amphibian with a different fertilization strategy: the axolotl, *A. mexicanum*. We therefore probed for zinc release during activation of immature oocytes which are more readily isolated from this internal fertilizer. Using confocal microscopy of *A. mexicanum* oocytes in FluoZin-3, we observed zinc release with ionomycin application from 5 of 6 oocytes (Fig 2A). Application of higher concentrations of ionomycin were required to activate zinc release from immature *A. mexicanum* and *X. laevis* oocytes (S3 Fig).

**Figure 2.**
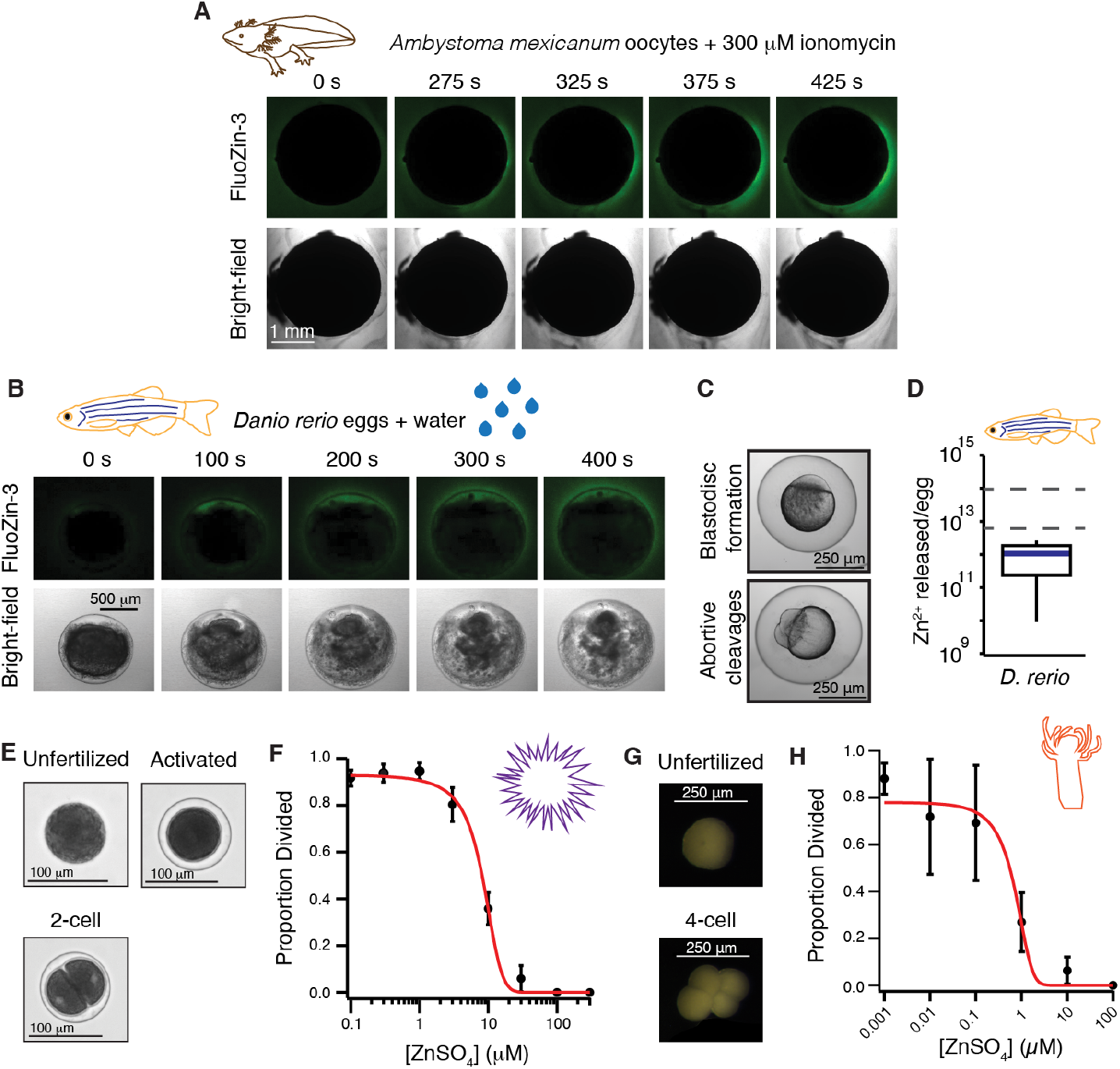
Zinc release and protection of eggs is shared in external fertilizers. (A) Representative FluoZin-3 and bright-field images of *A. mexicanum* oocytes before and after activation with 300 μM ionomycin (N=6 oocytes, 2 trials). (B) Representative images of FluoZin-3 and bright-field of *D. rerio* eggs releasing zinc upon activation with water (N=7 eggs, 4 trials). Eggs were in 50 μM FluoZin-3 (A & B). (C) Appearance of blastodisc and abortive cleavages confirmed activation for *D. rerio* eggs. (D) Box plot of zinc released per *D. rerio* egg upon activation as detected by fluorometry using FluoZin-3 (N=59-138 eggs, 5 trials). Dashed lines denote the 25^th^ and 75^th^ percentile of the data distribution for zinc released from *X. laevis* eggs following fertilization. (E) Representative images of *S. purpuratus* eggs and embryos. (F) Development was blocked in a concentration-dependent manner for *S. purpuratus* eggs in ZnSO_4_ (N=120-684 eggs, 5 trials). (G) Representative images of *H. symbiolongicarpus* unfertilized and divided eggs. (H) Development was blocked in a concentration-dependent manner for *H. symbiolongicarpus* eggs inseminated in extracellular ZnSO_4_ (N=143-1463 eggs, 5-6 trials). Errors are s.e.m.

Next, we expand our analyses to the more distantly related zebrafish, *D. rerio*, in which both activation of embryonic development and release of cortical granules from the egg are independent of fertilization. *D. rerio* eggs are activated during spawning in water, which typically coincides with mating but is independent of sperm entry [20, 31]. In the presence of FluoZin-3, we observed zinc release from *D. rerio* eggs upon hydration in 6 of 7 eggs (Fig 2B, Movie 4). Successful activation of cohort eggs was confirmed by blastodisc formation and the appearance of abortive cleavages in all eggs assayed (Fig 2C). FluoZin-3 fluorometry revealed that *D. rerio* eggs release an average of 1.1 ± 0.4 x 10^12^ zinc molecules upon activation (Fig 2D, Table 1). Together, these results suggest that zinc release upon activation may be conserved amongst vertebrate eggs.

To explore whether zinc-inhibition of fertilization and early embryonic development is shared with invertebrates, we fertilized eggs from the sea urchin *Strongylocentrotus purpuratus*, and the cnidarian *Hydractinia symbiolongicarpus*, in varying concentration of ZnSO_4_. For both species, embryonic development was blocked in a concentration-dependent manner, as assessed by the appearance of cleavage furrows (Fig 2E-H, S1 Table). To demonstrate that zinc effects on fertilization were not due to toxicity in embryos, fertilized *H. symbiolongicarpus* eggs were treated with 50 μM ZnSO_4_. We found that with zinc treatment 30 min after insemination, nearly all fertilized eggs developed to the larval stage. Together, these results are consistent with the hypothesis that zinc can inhibit fertilization and development in both vertebrate and invertebrate eggs.

To uncover the mechanisms by which zinc disrupts fertilization, we returned to the *X. laevis* model to discriminate whether zinc targets eggs or sperm. To do so, we assessed whether pre-treating eggs before fertilization with zinc was sufficient to block fertilization. For these experiments, *X. laevis* eggs were incubated with or without added zinc, then transferred to different solutions for insemination and embryonic development. Eggs pretreated with zinc failed to develop cleavage furrows, even when inseminated in a solution with no added zinc (Fig 3A). By contrast, eggs never exposed to zinc developed normally. Evidently, zinc altered the egg to interfere with fertilization and the earliest events of embryonic development. However, these data do not exclude the possibility that zinc also targets sperm to block fertilization.

**Figure 3.**
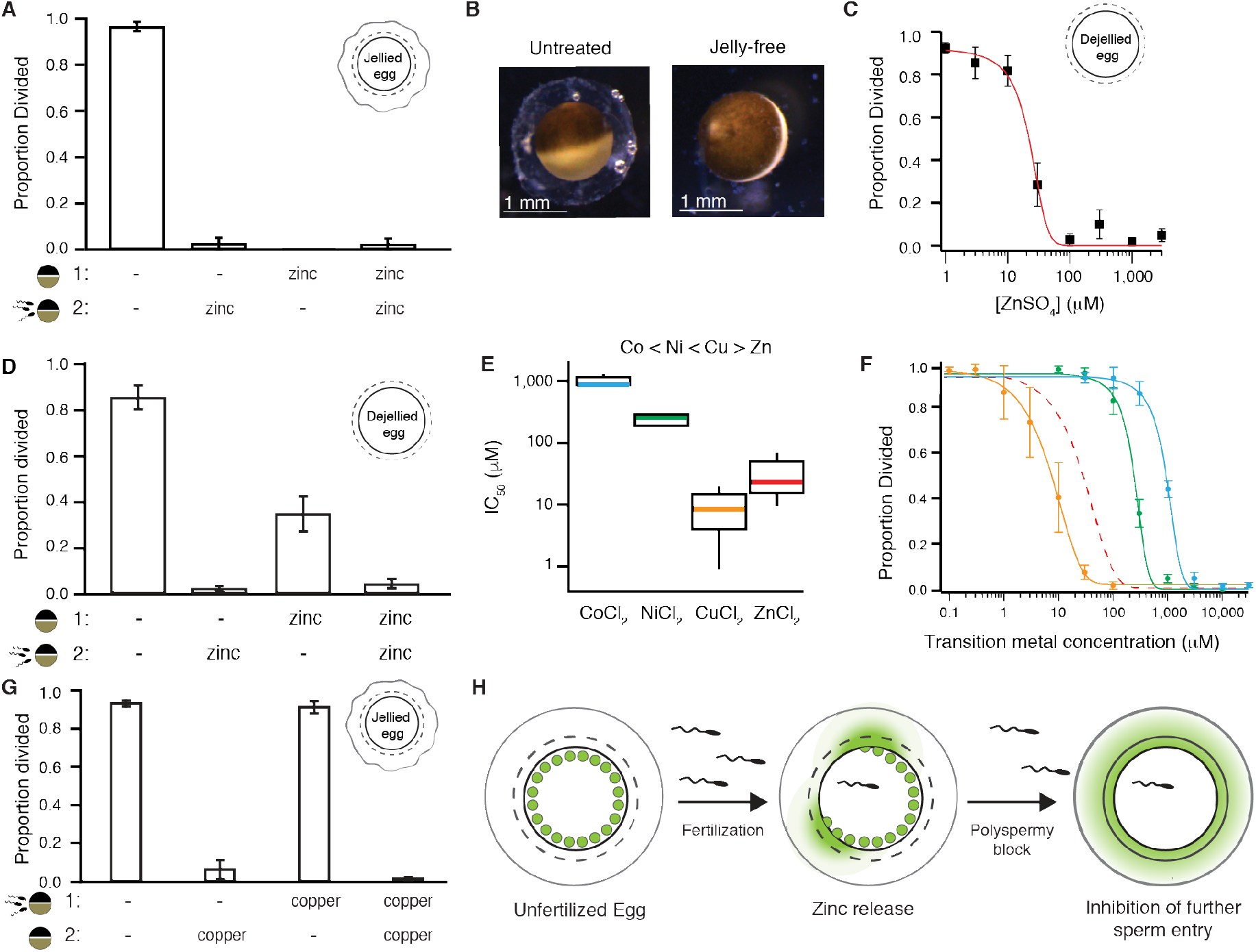
Zinc inhibits embryonic development by binding to proteins surrounding the egg. (A) Incidence of development of eggs pre-treated in ZnSO_4_ prior to insemination (N=34-150 eggs, 5 trials). (B) *X. laevis* eggs before and after jelly removal. (C) Incidence of development of jelly-free eggs in varying ZnSO_4_ concentrations (N=53-96 eggs, 6 trials). (D) Average proportion of cleavage furrow development from jellied eggs pre-treated in solution (*1*), with either 0 or 300 μM ZnSO_4_. After a 15 min treatment, eggs were then washed and moved to solution (*2*) for sperm addition with 0 or 300 μM TPEN (N=35-192, 5 trials). (E) Box plot distributions of the IC_50_s of inhibition of appearance of cleavage furrows in *X. laevis* eggs inseminated in cobalt (blue), nickel (green), copper (orange), or zinc (red). (F) Proportion of division of *X. laevis* eggs inseminated in varying concentrations of extracellular cobalt (N=148-295 eggs, 5 trials), nickel (N=136-283 eggs, 5 trials), or copper (N=230-321 eggs, 5 trials). (G) Incidence of cleavage furrow development from eggs inseminated in solution (*1*), with 0 or 100 μM CuCl_2_ then transferred to solution (*2*), with 0 or 100 μM CuCl_2_, 30 min after sperm addition (N=205-302 eggs, 5 trials). (H) Fertilization stimulates the release of zinc from cortical granules in a wave pattern to modify the envelope and jelly coat to inhibit sperm entry in *X. laevis*. All errors are s.e.m.

We then focused on the matrices surrounding the egg to probe how zinc disrupted fertilization. *X. laevis* eggs are surrounded by two distinct structures: the jelly coat and the envelope. The envelope is immediately outside of the plasma membrane and is comprised of five glycoproteins homologous to zona pellucida proteins that surround mammalian eggs [32–34]. Outside of the envelope is the jelly coat (Fig 3B), comprised of glycoproteins and enriched with salts [35–41].

To test whether the zinc-mediated disruption of fertilization required the jelly, we removed the jelly layer from *X. laevis* eggs and assayed for the appearance of cleavage furrows after insemination in varying concentrations of zinc. Even without the jelly, zinc prevented the appearance of cleavage furrows with a similar concentration-response relationship to eggs with an intact jelly layer (Figs 1E & 3C).

Although not required, the jelly may still play an important role in the zinc block of *X. laevis* fertilization. The jelly is known to bind to and buffer other metals [40] and may trap zinc near the egg. We assayed for the appearance of cleavage furrows in jelly-free *X. laevis* eggs pretreated with or without zinc and then inseminated in a second condition with or without zinc (Fig 3D). We predicted that if the jelly buffers zinc, then jelly-free eggs would develop cleavage furrows when inseminated after zinc pretreatment. Indeed, a third of the jelly-free eggs pretreated with zinc developed cleavage furrows following insemination with no added zinc. These results support a zinc-buffering role for the jelly, maintaining a high local zinc concentration, following the slow block.

The zinc disruption of *X. laevis* fertilization is reversible by chelation (Fig 1H), consistent with the hypothesis that zinc acts by binding to a protein or receptor and not by mediating a chemical reaction (*e.g*. reduction or oxidation). If extracellular zinc prevented fertilization through protein coordination, we would expect other transition metals would act similarly. Metal-protein interactions become more potent in a predictable pattern called the Irving-Williams series, which depicts the strength of divalent transition metal-protein complexes: Co < Ni < Cu > Zn [42]. To uncover how other transition metals in this series altered *X. laevis* fertilization, we inseminated in varying concentrations of extracellular copper, cobalt, or nickel. Development was blocked in a concentration-dependent manner in the presence of all transition metals tested (Fig 3E-F). Using the half-maximal concentration of each metal that blocks embryonic development, we determined that the potency of the transition metals indeed followed the Irving-Williams series [42]: copper acted with the highest affinity and cobalt with the lowest (Fig 3E, S1 Table). In follow-up experiments probing for possible toxic effects of these metals, we observed similar proportions of cleavage furrow development between eggs inseminated under control conditions and transferred to solutions with 0 or 100 μM CuCl_2_ 30 minutes following sperm addition (Fig 3G). We propose that release of zinc from the cortical granules upon fertilization modifies extracellular proteins to block entry by additional sperm (Fig 3H).

In conclusion, we report that zinc release upon fertilization or activation is shared amongst eggs from diverse vertebrates including mammals, amphibians, and teleost fish. Moreover, zinc is released by eggs from internal and external fertilizers from diverse phyla, as well as in species with varied mechanisms of fertilization and egg activation. We have also shown that early development is blocked in the presence of zinc in species from the phyla Cnidaria, Echinodermata, and Chordata. However, it has not yet been shown whether zinc release at fertilization occurs in invertebrate eggs. Our observations are consistent with the hypothesis that zinc is derived from cortical granules released during the slow block to polyspermy and suggest that this extracellular zinc transforms the egg to prevent multiple fertilizations. Although the details on how zinc prevents sperm entry are yet to be determined, we speculate that extracellular zinc may modify the envelope to stop sperm entry, or target sperm to interfere with fertilization, or a combination of the two. Taken together, these data suggest that extracellular zinc protection of eggs from multiple fertilizations is shared by diverse species separated by hundreds of millions of years of evolution.

## Methods

### Ethics

All vertebrate animal procedures were conducted using accepted standards of humane animal care and were approved by the Animal Care and Use Committee at the University of Pittsburgh.

### Animals

*Xenopus laevis* (frog) adults were obtained commercially (NASCO, Fort Atkinson, WI), as were *Ambystoma mexicanum* (axolotl) adults (Ambystoma Genetic Stock Center, Lexington, KY) and housed separately at 18°C with a 12/12-hour light/dark cycle. *Danio rerio* (zebrafish) adults (5-17 months) were lab bred TU-AB strain and housed at 27°C with a 14/10-hour light/dark cycle.

Some *Strongylocentrotus purpuratus* (purple sea urchin) gametes were a generous gift from Veronica Hinman and some were obtained from commercially purchased adults (Marinus Scientific, Long Beach CA) and housed at 15°C with a 12/12-hour light/dark cycle. *Hydractinia symbiolongicarpus* (cnidaria) were sexually mature, lab bred colonies (MN291-10 and MN295-8, male and female, respectively) grown on glass microscope slides and housed at 22-23°C with an 8/16-hour light/dark cycle.

### Reagents

1 M MgSO_4_ solution and 0.1 M ZnCl_2_ solution were purchased from Sigma-Aldrich (St. Louis, MO). TPEN was purchased from Tocris (Bristol, United Kingdom) and human chorionic gonadotropin (hCG) was purchased from Henry Schien (Melville, NY). Unless noted otherwise, all materials were purchased from Thermo Fisher Scientific (Waltham, MA).

### Solutions

Modified Ringers (MR) solution was used for *X. laevis* fertilization experiments. MR contains (in mM): 100 NaCl, 1.8 KCl, 2.0 CaCl_2_, 1.0 MgCl_2_, and 5.0 HEPES, pH 7.8, and is filtered using a sterile, 0.2 μm polystyrene filter [43]. Embryonic development assays were performed in 33% MR diluted in DDH_2_O (MR/3). Various chemicals were added to MR/3, which contained final concentrations of <0.5% DMSO or ethanol.

Oocyte Ringers 2 (OR2) solution was used to rinse *X. laevis* and *A. mexicanum* oocytes after collagenase treatment. OR2 is comprised of (in mM): 82.5 NaCl, 2.5 KCl, 1 MgCl_2_, and 5 mM HEPES, pH 7.6, and is filtered using a sterile, 0.2 μm polystyrene filter [44].

ND96 was used to store *X. laevis* and *A. mexicanum* oocytes. ND96 is comprised of (in mM): 96 NaCl, 2 KCl, 1 MgCl_2_, 10 HEPES, pyruvic acid, and 10 mg/L gentamycin at pH 7.6 and is filtered with a sterile 0.2 μm polystyrene filter.

Oocyte culture media (OCM) was used for *in vitro* maturation of *X. laevis* oocytes. OCM is comprised of: 40% water, 60% L-15 media, 60 μg/mL gentamycin, and 0.4 mg/mL BSA [45].

Lab-made artificial sea water (spASW) was used for *S. purpuratus* development assays. The spASW is comprised of (in mM): 470 NaCl, 10 KCl, 11 CaCl_2_, 29 MgSO_4_, 27 MgCl_2_, and 2.5 NaHCO_3_, pH 8, and is filtered using a sterile, 0.2 μm polystyrene filter [46].

Commercial artificial sea water (hsASW) was used for *H. symbiolongicarpus* development assays. The hsASW is comprised of solubilized Instant Ocean Reef Crystals (Instant Ocean Spectrum Brands) at 28 parts per thousand.

### Collection of gametes

*X. laevis and A. mexicanum oocytes:* Oocytes were collected from ovarian sacs obtained from *X. laevis* females anesthetized with a 30-min immersion in 1.0 g/liter tricaine-S (MS-222), pH 7.4 or from *A. mexicanum* females euthanized by a 30-minute immersion in 3.6 g/L tricaine-S at pH 7.4. Following excision, ovarian sacs were manually pulled apart, then dispersed by a 90-minute incubation in ND96 supplemented with 1 mg/ml collagenase. Collagenase was removed by repeated washes with OR2, and healthy oocytes were sorted and stored at 14°C in ND96 with sodium pyruvate and gentamycin.

*X. laevis sperm and eggs:* Eggs were collected from sexually mature females. Egg laying was stimulated by injection with 1,000 IU of hCG into their dorsal lymph sac. Following injection, females were housed overnight for 12-16 hours at 14-16°C. Typically, egg-laying began within 2 hours of moving to room temperature. Eggs were collected on dry petri dishes and used within 10 minutes of being laid.

Sperm were obtained from testes harvested from sexually mature *X. laevis* males [3, 4, 40]. Following euthanasia by a 30-minute immersion in 3.6 g/L tricaine-S (pH 7.4), testes were dissected and cleaned by manual removal of residual fat and vasculature. Cleaned testes were stored at 4°C in MR for use on the day of dissection or in L-15 medium for use up to one week later. Sperm were extracted by mincing 1/10 of a testis in 200-500 μL of MR and were used within 1 hr of collection.

For experiments sequentially treating eggs with different conditions, eggs were incubated in an initial experimental solution, washed three times by moving between petri dishes containing the final treatment using plastic transfer pipettes, and then placed in the final treatment for experimental observation. Two types of these sequential treatment assays are reported here: transfer before insemination (*e.g*. Fig 1H) and transfer after insemination (*e.g*. Fig 1I). When transferred between treatments before fertilization, eggs were incubated in the starting solution for 15 minutes and inseminated in the transfer solution. When transferred after insemination, eggs and sperm were incubated together in the starting solution for 30 minutes, then transferred.

*D. rerio: D. rerio* gametes were obtained from mating pairs selected at random. To stimulate spawning, mating pairs were housed in the same tank overnight, separated by a divider [20, 47]. In the morning, the females were anesthetized in 0.25 mg/mL MS-222 (pH 7.2), rinsed in DDH_2_O, and patted dry. Cohorts of eggs were retrieved from anesthetized females by applying pressure to their abdomens [20]. Dry eggs were collected with a 200 μL pipette tip and moved to a dry imaging slide.

*S. purpuratus:* Gametes were collected from spawning *S. purpuratus* adults. Spawning was induced by manual agitation or injection with 100-500 μL of 0.5 M KCl followed by agitation [46]. Sperm were collected directly from the animal using a 10 μL pipette and then transferred into a 1.7 ml capped tube. Eggs were collected following release into a beaker containing spASW, then filtered through a 100 μm filter.

*H. symbiolongicarpus: H. symbiolongicarpus* gametes were collected from spawning adults. Upon the first light exposure for the day, spawning was induced following separation of male and female colonies [48]. Gametes were released within 60-90 minutes of light exposure. Eggs were collected from the water surrounding spawning females, filtered through a 20 μm strainer, and maintained in hsASW. Sperm were collected from the water surrounding spawning males using a 1 mL pipette and maintained in hsASW.

### Confocal microscopy of extracellular zinc during fertilization or activation

Zinc release from *X. laevis, D. rerio*, and *A. mexicanum* gametes was imaged using FluoZin-3 and a TCS SP5 confocal microscope (Leica Microsystems, Wetzlar, Germany) equipped with a Leica 506224 5X objective. FluoZin-3 was excited with a 488 nm visible laser, and the emission between 500-600 nm was collected. Using a galvo scanner with unidirectional (600 Hz) scanning, FluoZin-3 and bright-field images were taken every 3-5 seconds for up to 25 minutes with a depth of 3 μM, the laser strength at 30%, and gain at 10%. Images were analyzed using LAS AF (version 3.0.0 build 834) and ImageJ [49] software packages.

*X. laevis*: To image extracellular zinc during *X. laevis* fertilization, sperm were added to dejellied eggs bathed in MR/3 with 50 μM FluoZin-3. Sperm prepared in MR/3, was pipetted near the eggs 1 minute after image acquisition had begun. Control experiments used the same experimental design with 1.5 mM TPEN. A similar experimental design was employed to image extracellular zinc during activation where *X. laevis* eggs or oocytes were activated with 10 or 200 μM ionomycin treatment respectively in MR/3 with no sperm addition.

*A. Mexicanum:* Extracellular zinc was imagined during *A. mexicanum* oocyte activation by incubating these cells with 50 μM FluoZin-3 in MR/3. These oocytes were activated with 300 μM ionomycin application, 1 min after imaging began.

*D. rerio*: To image extracellular zinc during *D. rerio* egg activation, dry eggs were placed on a microscope slide. Eggs were activated by application of 50 μM FluoZin-3 in DDH_2_O, 1 minute after image acquisition had begun.

### Bright-field microscopy

*X. laevis:* A stereoscope (Leica Microsystems, Wetzlar, Germany) equipped with a Leica 10447157 1X objective and DFC310 FX camera was used to image *X. laevis* eggs and embryos. Images were analyzed using LAS (version 3.6.0 build 488) software and Photoshop (Adobe). For the jelly removal assay, *X. laevis* eggs were imaged using an Edmund Optics stereomicroscope with a 10x objective, fitted with a pixiLINK digital camera and the μScope Essential 64x software pixiLINK, Canada). The diameter of the egg and the surrounding jelly coat were determined in Adobe Illustrator (San Jose, CA).

*D. rerio* egg & *S. purpuratus:* Eggs and embryos from *D. rerio* and *S. purpuratus* were imaged on an inverted Olympus IX73 stereoscope equipped with an Olympus UPlanFL N 10X objective, Olympus TL4 light source, and Olympus U-LS30-3 camera. Images were analyzed using Photoshop (Adobe).

*H. symbiolongicarpus* eggs and embryos were imaged using Zeiss Discovery.V20 stereoscope equipped with a Lumenera Infinity3s camera and Zeiss KL1500 LCD light source. Images were analyzed using Photoshop (Adobe).

### *In vitro* maturation of *X. laevis* oocytes

*X. laevis* oocytes arrested in prophase I were matured into metaphase II arrested oocytes by incubating immature oocytes in OCM supplemented with 8 μM progesterone, at 18°C for 12-14 hours. Maturation was visually confirmed by the appearance of a maturation spot.

### Fertilization and embryonic development assays

*X. laevis:* For each experimental trial, development of *X. laevis* embryos was assessed from approximately 20-40 eggs in each experimental condition. 20-90 μL of the sperm suspension was used to fertilize eggs depending on the volume of the dish. Approximately 90-120 minutes after insemination, the appearance of cleavage furrows was used to assess the initiation of embryonic development. In the case of the TPEN timing assay, eggs were imaged every 10 minutes beginning at 60 minutes post insemination to assess for development to the 2-cell stage. Each assay was repeated at least three times with gametes from different males and females and on different experiment days.

*S. purpuratus: S. purpuratus* sperm were activated by 5:1,000 dilution into spASW. 30 μL of activated sperm were added to a 4 ml suspension eggs in spASW supplemented with varying concentration of ZnSO_4_. Successful fertilization was visually confirmed 2 minutes after sperm addition by the raising of the fertilization envelope. Development was assayed 90-120 minutes post-fertilization based on the appearance of cleavage furrows.

*H. symbiolongicarpus:* For development assays, equal volumes of *H. symbiolongicarpus* sperm and egg solutions were mixed together with varying concentrations of ZnSO_4_ in hASW. Development was assayed at 60 minutes post-fertilization based on the appearance of cleavage furrows.

### *H. symbiolongicarpus* viability assay

To assay for possible toxic side-effects of zinc on *H. symbiolongicarpus* embryos, equal volumes of *H. symbiolongicarpus* sperm and egg solutions were mixed together in an hsASW solution lacking zinc. 30 minutes following insemination, eggs were transferred with a plastic transfer pipette to a 50mL Falcon tube. Approximately 20 mL of hsASW with 10μM ZnSO_4_ was added. Eggs were placed in a petri dish, and the process repeated until 50 mL of hsASW + ZnSO_4_ was used. Eggs were finally placed in a petri dish in the hsASW + ZnSO_4_ solution and incubated 2-3 days at room temperature. Proportion of living larvae was assessed the following day.

### Removing the jelly coat surrounding *X. laevis* eggs

In some experiments, the jelly coat surrounding *X. laevis* eggs was removed prior to fertilization [40, 43]. To do so, eggs were agitated in 45 mM β-mercaptoethanol in MR/3 (pH 8.5) for 2-3 minutes. Once removal of the jelly coat was confirmed by visual observation, eggs were neutralized in MR/3 (pH 6.5), followed by three washes in MR/3 (pH 7.8).

### Parthenogenic egg activation

*X. laevis*: For development assays following parthenogenic activation, eggs were placed in 10 μM ionomycin for 7 minutes, washed in MR/3 three times, and incubated in MR/3 for 120-150 minutes before developmental assessment.

*A. mexicanum*: Activation was induced by incubation of *A. mexicanum* eggs in MR/3 and incrementally adding 3 mM ionomycin to the solution, accumulating to a final concentration of ~300 μM ionomycin.

*D. rerio:* Activation was induced by hydrating the eggs with DDH_2_O[20]. Successful egg activation was assayed based on the appearance of a blastodisc and abortive cleavages, occurring 90- and 120-min post-hydration, respectively.

### FluoZin-3 Fluorometry

*X. laevis*: To quantify extracellular zinc released during fertilization or parthenogenic activation, jelly was removed from batches of 30-100 freshly ovulated eggs. Jelly-free eggs were then inseminated with sperm or activated with 10 μM ionomycin. The solution (MR/3) surrounding the eggs was collected 45 minutes after sperm addition or 30 minutes after ionomycin addition. The fertilization solutions were then sedimented at 3000 rpm for 5 minutes to pellet sperm, and the supernatant was transferred to a new tube.

*D. rerio:* To quantify extracellular zinc released during activation of *D. rerio* eggs, batches of 40-120 freshly ovulated eggs were parthenogenically activated in DDH_2_O. 45 min following DDH_2_O hydration, the solution surrounding the eggs was collected and mixed with equal parts DDH_2_O and MR to create an MR/3 solution. The zinc content of each MR/3 sample was quantified using FluoZin-3 photometry.

*X. laevis and D. rerio:* FluoZin-3 tetrapotassium salt was dispensed from a 1 mM stock in water for a final experimental concentration of 60 nM. Fluorescence intensity measurements were recorded in a 1 mm quartz cuvette, in a Fluorolog3 spectrophotometer with FluoEssence software (both from HORIBA, Jobin Yovon). FluoZin-3 containing samples were excited with 492 nm light, and emission was recorded at 514 nm with 3 nm slit widths. The raw photometric signals were corrected for by subtracting the FluoZin-3 free background, collected prior to adding FluoZin-3 to each sample. The zinc was quantified using a standard curve[40, 50, 51] calculated with the following equation:

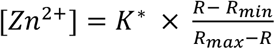

where the constants R_min_ (1 nM), R_max_ (100 nM), and K* were obtained from MR/3 supplemented with known amounts of ZnSO_4_ ranging from 100 pM to 1 μM fitted to a Hill equation [40, 50]. Although there was little variability between experiments, standard curves were generated for each experimental trial using the exact solution used for fertilization or activation. The following negative controls were assayed, each with no substantial zinc signal: MR/3 alone, MR/3 with sperm alone (sperm were removed by sedimentation prior to sample measurements), eggs alone (no sperm or ionomycin added), and MR/3 with ionomycin but no cells.

To determine the number of zinc ions released with fertilization or parthenogenic activation, the total zinc concentration measured by FluoZin-3 photometry was multiple by the dilution factor for that sample and Avogadro’s number, and divided by the total volume of solution in which eggs were inseminated from each trial and the number of eggs per trial.

*X. laevis* eggs reportedly contain 65.8 ± 4 ng/egg of zinc [30]. The number of zinc ions contained by each *X. laevis* egg was calculated by dividing the zinc mass by the atomic mass of zinc (65.38), then multiplying by Avogadro’s number.

### Inductively coupled plasma mass spectrometry (ICP-MS)

Eggs were dejellied and transferred to a clean 35 mm petri dish containing MR/3. All transfers were performed with disposable glass transfer pipettes. Eggs were activated with 100 nM-10 μM ionomycin (free acid) application and incubated at room temperature 30 minutes. The incubation solution was then collected and transferred to 15 mL conical tubes for storage at −20°C with up to 2.5% nitric acid prior to analysis by ICP-MS. Eggs were transferred to fresh MR/3 to assess activation 60-90 minutes after ionomycin addition. Biological controls included: eggs in MR/3 (no ionomycin added), MR/3 alone, and MR/3 with 100 nM ionomycin.

Released zinc was quantified with a PerkinElmer NexION 300X ICP-MS. On each experimental day, the instrument was calibrated with a five-point calibration curve. A blank consisting of 2% sub-boil distilled trace metal grade nitric acid was run every 7-10 samples to rule out signal memory effects. Reported values reflect the zinc concentration from the number of activated eggs, following subtraction of the solution background (obtained from the egg-free MR/3 with ionomycin control).

### Proteomic and RNA-sequencing (RNA-seq) analysis

Zinc-transporters expressed in *X. laevis* eggs were identified by interrogating proteomic [28] and RNA-seq [29] datasets for SLC30 or SLC39 gene names.

### Data availability

The authors declare that all relevant data are included in this paper and the supplementary information files.

## Acknowledgements

We thank Z. Crowell for technical assistance, V. Hinman for help troubleshooting, and K. Komondor for critical review of the manuscript. This work was supported by a Margaret A. Oweida Predoctoral Fellowship to K.L.W., an American Heart Association Predoctoral Fellowship 18PRE33960391 to M.T., a National Science Foundation grant 1557339 to M.L.N, and National Institute for Health grants R01GM125638 and R00HD069410 to A.E.C. Undergraduate summer research fellowships from Howard Hughes Institute Science Education Grant #52008122 provided summer stipends for M.E.C., M.B.L., S.P., M.L.S., and B.W.W.

## Author Contributions

**Conceptualization:** Katherine L. Wozniak, Rachel E. Bainbridge, Maiwase Tembo, Anne E. Carlson

**Formal Analysis:** Katherine L. Wozniak, Rachel E. Bainbridge, Anne E. Carlson

**Funding Acquisition:** Katherine L. Wozniak, Maiwase Tembo, Anne E. Carlson

**Investigation:** Katherine L. Wozniak, Rachel E. Bainbridge, Dominique W. Summerville, Maiwase Tembo, Wesley A. Phelps, Monica L. Sauer, Bennett W. Wisner, Madelyn E. Czekalski, Srikavya Pasumathy, Meghan L. Hanson, Melania B. Linderman, Catherine H. Luu, Madison E. Boehm, Daniel J. Bain, Anne E. Carlson

**Methodology:** Katherine L. Wozniak, Rachel E. Bainbridge, Dominique W. Summerville, Maiwase Tembo, Wesley A. Phelps, Monica L. Sauer, Bennett W. Wisner, Madelyn E. Czekalski, Srikavya Pasumathy, Miler T. Lee, Matthew L. Nicotra, Daniel J. Bain, Katherine M. Buckley, Steven M. Sanders, Anne E. Carlson

**Project administration:** Katherine L. Wozniak, Rachel E. Bainbridge, Anne E. Carlson

**Resources:** Anne E. Carlson, Miler T. Lee, Matthew L. Nicotra, Daniel J. Bain, Katherine M. Buckely, Stephen M. Sanders

**Supervision:** Anne E. Carlson, Miler T. Lee, Daniel J. Bain, Matthew L. Nicotra, Katherine M. Buckley

**Visualization:** Katherine L. Wozniak, Rachel E. Bainbridge, Maiwase Tembo, Anne E. Carlson

**Writing – original draft:** Katherine L. Wozniak, Rachel E. Bainbridge, Anne E. Carlson

**Writing – Reviewing and editing:** Katherine L. Wozniak, Rachel E. Bainbridge, Dominique W. Summerville, Maiwase Tembo, Wesley A. Phelps, Monica L. Sauer, Bennett W. Wisner, Madelyn E. Czekalski, Steven M. Sanders, Katherine M. Buckley, Daniel J. Bain, Matthew L. Nicotra, Miler T. Lee, Anne E. Carlson

**Competing Interests:** The authors have declared that no competing interests exist.

## Supporting information

Supplementary Figures 1–3

Supplementary Table 1

Movies 1-4

## Supplementary Materials

### Supplementary Figures

**Figure S1.**
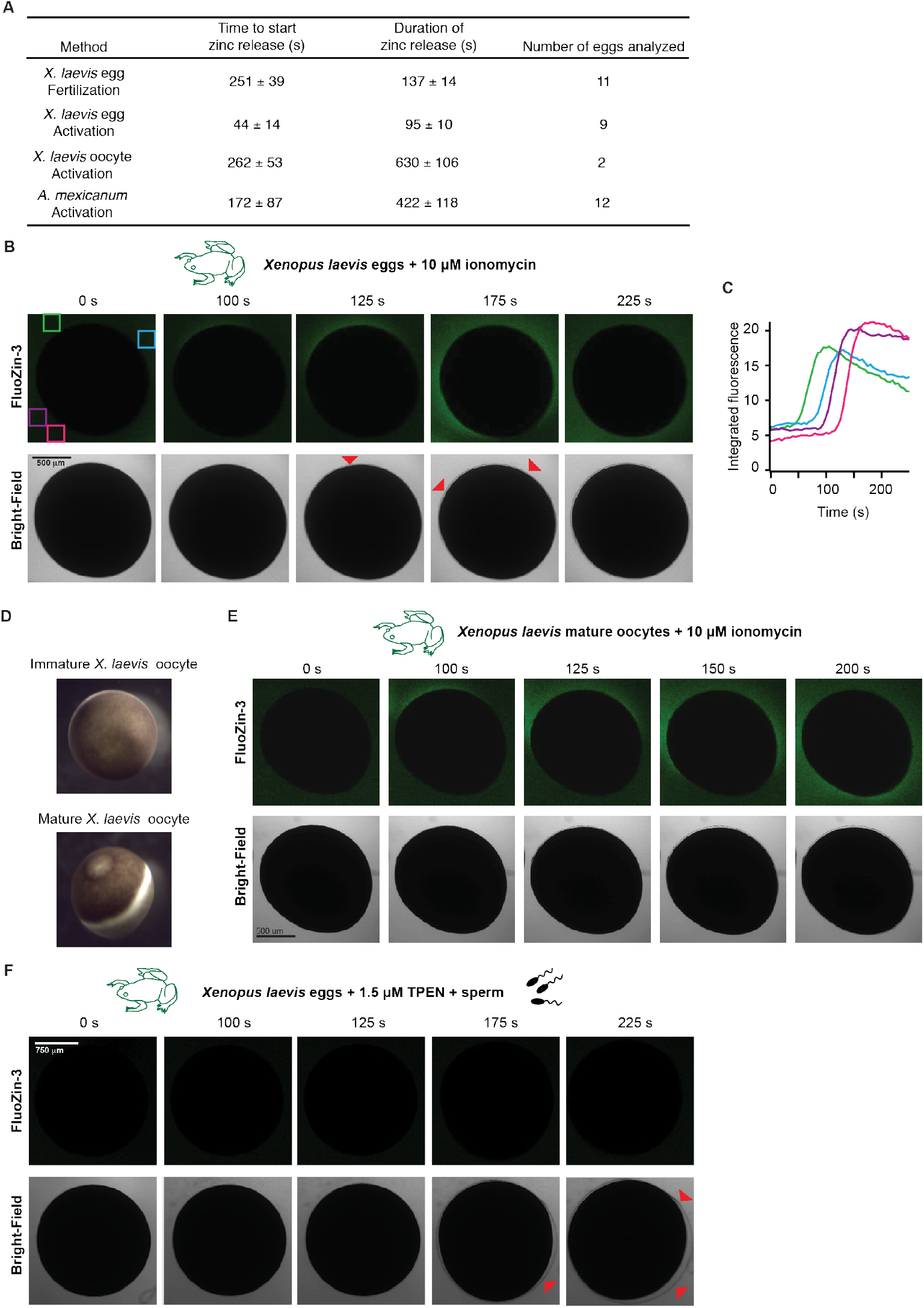
Activation in the absence of fertilization promotes zinc release from *X. laevis* eggs and oocytes. (A) Kinetics of zinc release in activation of *X. laevis* and *A. mexicanum* eggs and oocytes. (B) Parthenogenic activation of *X. laevis* eggs (N=9 eggs, 5 trials) or *in vitro* matured oocytes (E; N=9 eggs, 3 trials) with 10 μM ionomycin in the presence of FluoZin-3 also induced zinc exocytosis. Red arrowheads highlight the lifting of the fertilization envelope (B, F). (C) Changes in FluoZin-3 fluorescence upon parthenogenic activation were detected by region of interest analysis. Integrated fluorescence relative to time of ionomycin addition, detected by region of interest analysis (indicated by colored boxes in upper left image). (D) Representative images of immature and *in vitro* matured oocytes. (F) Treatment of *X. laevis* eggs in FlouZin-3 with the zinc chelator TPEN abolished fertilization-induced zinc release (N=8 eggs, 2 trials).

**Figure S2.**
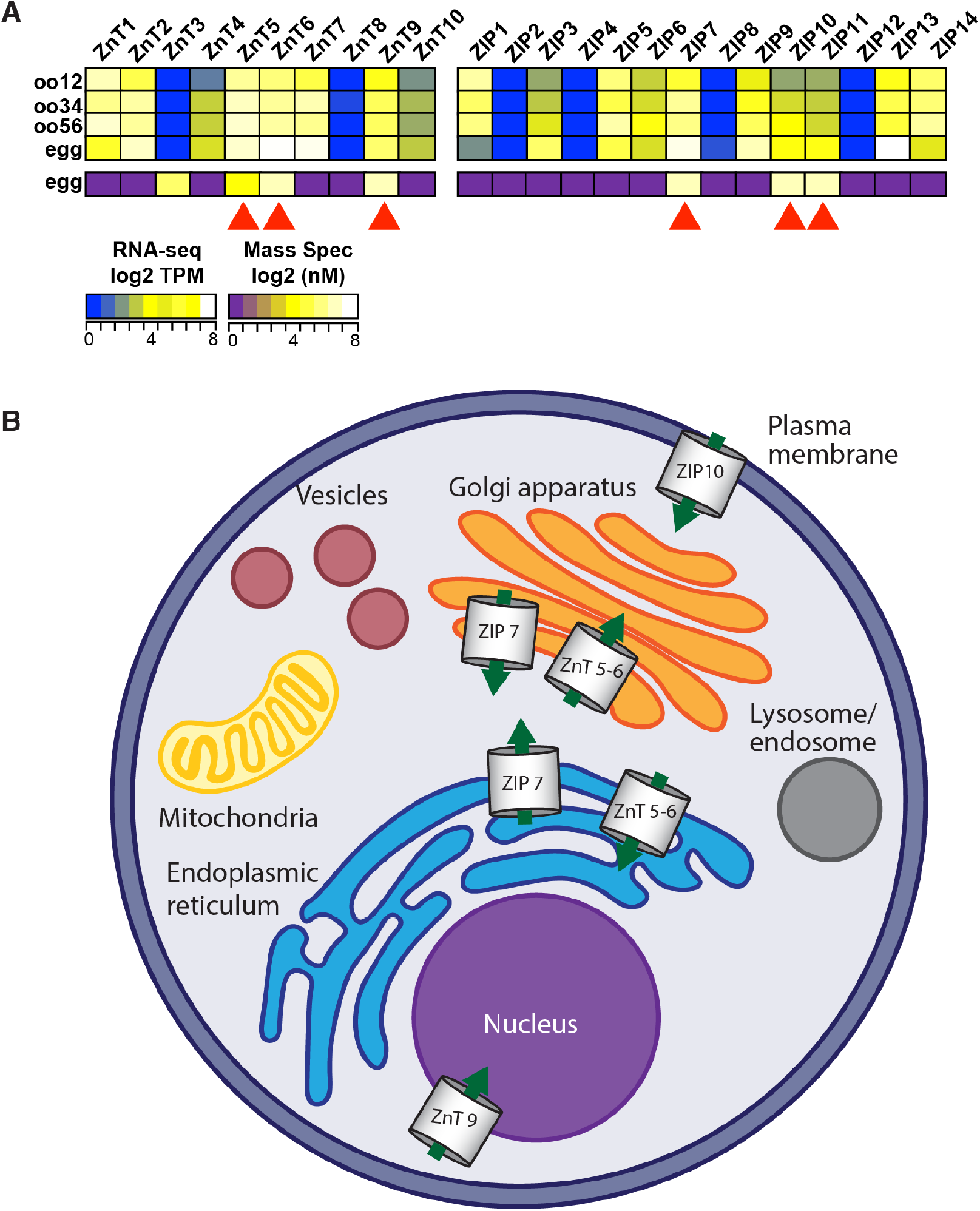
Zinc transporters found in the *X. laevis* female gametes. (A) Heatmaps of the expression levels of the two known families of zinc transporters ZnTs (encoded by the SLC30 gene family) and ZIPs (encoded by the SLC39 gene family) at the developmental stages indicated. Transcript levels (from [29]) as determined by RNA-seq-based transcriptomics study, depicted in log2 transcripts per million (TPM) (upper). Protein concentrations (from[28]) as determined by mass spectrometry-based proteomics study, depicted in log2 nanomolar (lower). Arrowheads highlight the six zinc transporters with both RNA and protein present in *X. laevis* eggs: ZnT5, ZnT6, ZnT9, ZIP7, ZIP10, and ZIP11. (B) Schematic of cellular localization of zinc transporters in the cell. ZnT5, ZnT6, and ZIP7 localize to the Golgi apparatus and the endoplasmic reticulum [52, 53] and ZnT9 to the nucleus [52, 53]. ZIP10 is localized to the plasma membrane and allows transport of extracellular zinc into the cytoplasm [52, 54]. Cellular localization of ZIP11 is not yet clear, though it is predicted to traffic to the nucleus and/or Golgi apparatus [52].

**Figure S3.**
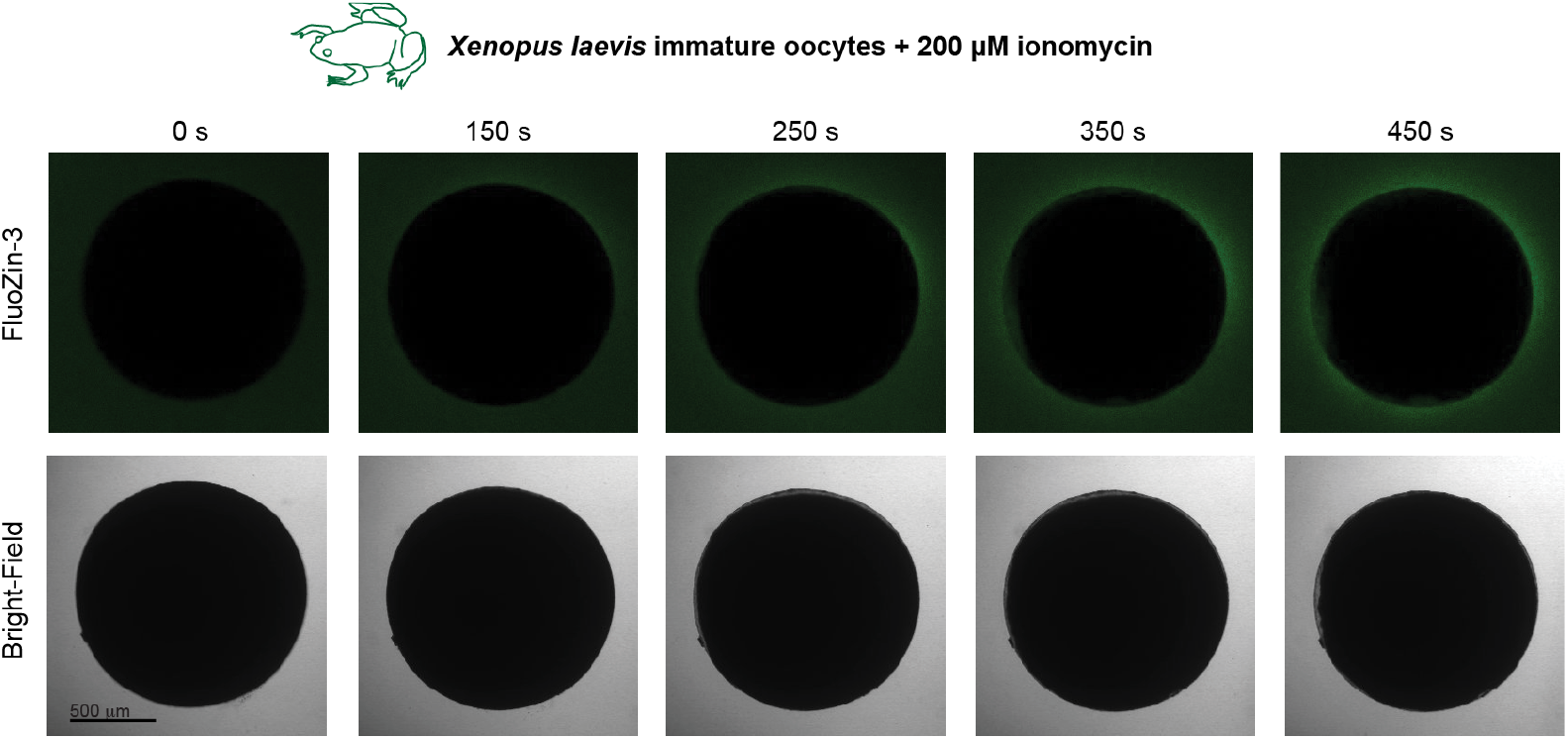
Activation of immature *X. laevis* eggs. Zinc released upon activation of immature *X. laevis* oocytes with 200 μM ionomycin (N=15 eggs, 7 trials).

**Table S1.** Concentration-response for eggs fertilized in various transition metals.

|  | *Xenopus laevis* | *Strongylocentrotus purpuratus* | *Hydractinia symbiolongicarpus* |
| --- | --- | --- | --- |
| ZnSO<sub>4</sub> | 31 ± 10 µM<br>169 - 269 eggs<br>7 trials | 9 ± 1 µM<br>120 - 684 eggs<br>5 trials | 4 ± 2 µM<br>143 - 1,463 eggs<br>5 - 6 trials |
| ZnSO<sub>4</sub><br>Jelly-free eggs | 23 ± 3 µM<br>53 - 96 eggs<br>6 trials | — | — |
| ZnCl<sub>2</sub> | 30 ± 8 µM<br>157 - 326 eggs<br>8 trials | — | — |
| MgSO<sub>4</sub> | No inhibition<br>182 - 383 eggs<br>5 trials | — | — |
| CuCl<sub>2</sub> | 9.1 ± 3.0 µM<br>230 - 321 eggs<br>5 trials | — | — |
| NiCl<sub>2</sub> | 224 ± 23 µM<br>136 - 283 eggs<br>5 trials | — | — |
| CoCl<sub>2</sub> | 971 ± 92 µM<br>148 - 295 eggs<br>5 trials | — | — |

**Movie 1.** Extracellular zinc imaged with 50 μM FluoZin-3 (left) and bright-field (right) confocal microscopy on *X. laevis* eggs, before and after sperm application. Time stamps are in minutes.

**Movie 2.** Extracellular zinc imaged with bright-field (left) and 50 μM FluoZin-3 (right) confocal microscopy on *X. laevis* eggs, before and after activation with 10 μM ionomycin addition. Time stamps are in minutes.

**Movie 3.** Bright-field (left) and 50 μM FluoZin-3 images of *X. laevis* eggs collected in the presence of 50 μM FluoZin-3 and 1 mM TPEN, before and after sperm addition.

**Movie 4.** Extracellular zinc imaged with 50 μM FluoZin-3 (left) and bright-field (right) confocal microscopy on *D. rerio* eggs, before and after activation by hydration with DDH_2_O. Time stamps are in minutes.

